# Sensitivity to food and cocaine cues are independent traits in a large sample of heterogeneous stock rats

**DOI:** 10.1101/2020.05.13.066944

**Authors:** Christopher P. King, Jordan A. Tripi, Alesa R. Hughson, Aidan P. Horvath, Alexander C. Lamparelli, Katie L. Holl, Apurva Chitre, Oksana Polesskaya, Jerry B. Richards, Leah C. Solberg Woods, Abraham A. Palmer, Terry E. Robinson, Shelly B. Flagel, Paul J. Meyer

## Abstract

Sensitivity to cocaine and its associated stimuli (“cues”) are important factors in the development and maintenance of addiction. Rodent studies suggest that this sensitivity is related, in part, to the propensity to attribute incentive salience to food cues, which, in turn, contributes to the maintenance of cocaine self-administration, and cue-induced relapse of drug-seeking. Whereas each of these traits has established links to drug use, the relatedness between the individual traits themselves has not been well characterized in preclinical models. To this end, the propensity to attribute incentive salience to a food cue was first assessed in a large population of 2716 outbred heterogeneous stock rats. We then determined whether this was associated with performance in two paradigms (cocaine conditioned cue preference and cocaine contextual conditioning). These measure the unconditioned locomotor effects of cocaine, as well as conditioned approach and the locomotor response to a cocaine-paired floor or context. There was large individual variability and sex differences among all traits, but they were largely independent of one another in both males and females. These findings suggest that these traits may contribute to drug-use via independent underlying neuropsychological processes.

## Introduction

Complex interactions between a host of genetic and environmental factors are thought to result in a number of intermediate traits that may confer vulnerability to develop impulse control disorders, including addiction. There has been considerable preclinical research on traits that predict drug self-administration behavior and susceptibility to relapse in order to better understand the neuropsychological bases of these vulnerability factors. For example, in rodents, behavioral phenotypes thought to influence drug-taking and -seeking behavior include the propensity to attribute incentive value to reward cues^1–3^, novelty-seeking^4–6^, locomotor response to novelty^7–9^, and impulsivity^10–13^.

Of these traits, we have been especially interested in how individual variation in the propensity to attribute incentive salience to rewards and their associated stimuli (“cues”) influence the development of addiction-like behavior. When delivery of a food reward is paired with presentation of a cue (conditioned stimulus, CS; usually a lever) some rats (sign-trackers, ST) come to approach and interact with the CS itself^14,15^, whereas during the CS period others (goal-trackers, GT) approach and interact with the food cup^15,16^. These phenotypic differences predict a number of addiction-related behaviors^17^, including responses to drug and drug cues^3,18^, the ability of drug cues to support drug-taking behavior^19^, and the ability of drug cues to motivate drug-seeking behavior^1,2,20^. Sign-tracking is also associated with other traits thought to confer vulnerability of addiction, most notably, impulsivity and poor top-down attentional control over behavior^21–23^. However, the extent to which sign-tracking is associated with other unconditioned or conditioned drug responses is not well understood.

We have begun to address sign-tracking in its relation to other traits as part of an ongoing genome-wide association study (GWAS), and report here initial results from a sample of 2,716 outbred heterogeneous stock (HS) rats. The HS rats originate from a cross of 8 inbred strains, now maintained as a unitary outbred population. The propensity to approach a food cue was measured in all HS rats using the Pavlovian conditioned approach (PavCA) procedure; we then determined the extent to which this was correlated with 1) The unconditioned immediate locomotor activating effects of cocaine^24^, 2) the conditioned approach response to a cocaine-paired floor stimulus^25^, and 3) the conditioned locomotor response to a cocaine-paired context^26^. We also assessed whether any of these effects were sex-dependent.

## Results

The tendency to attribute incentive salience to a food-CS was assessed in two cohorts of HS rats using an identical PavCA procedure, one at the University at Buffalo (UBuff; n = 1528) and another at the University of Michigan (UMich; n = 1188), although it should be noted that the timing of this test varied between the two cohorts (see **Table 1)**. Both sites then examined the reinforcing properties of the lever-CS during a conditioned reinforcement (CRF) procedure before separately measuring each cohort for either conditioned approach to towards a cocaine-paired floor, or conditioned locomotion in a cocaine-paired context. Here, we first present the PavCA data from the UBuff cohort replicating the findings of the UMich cohort **(Fig. 1, 2)** before examining the relationship with unconditioned and conditioned cocaine responses; the UMich cohort of rats have been described in detail in reference^27^.

**Figure 1:**
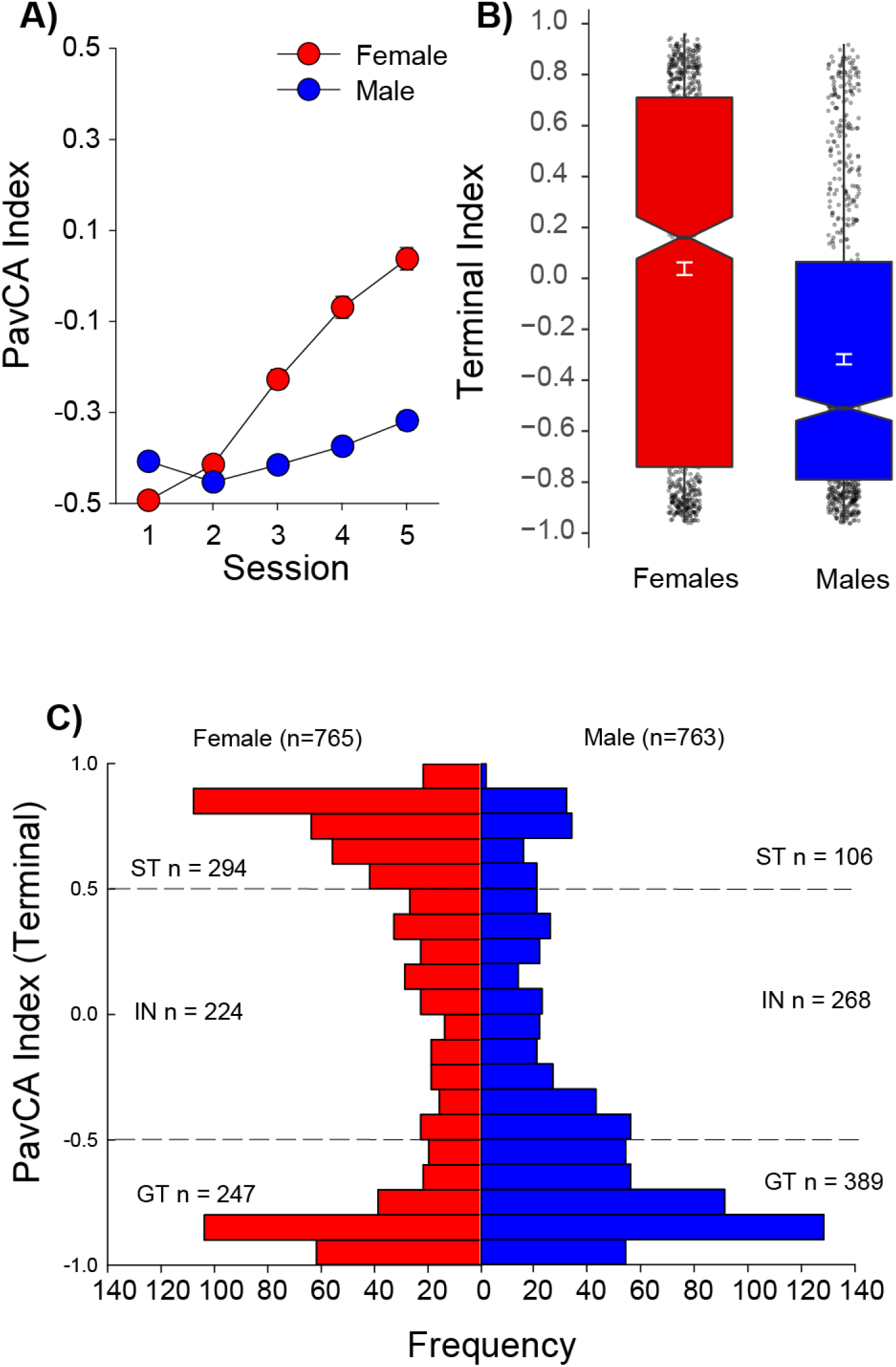
Sex differences during Pavlovian conditioned approach (UBuff). Across 5 sessions (total *n* = 1528), **A)** female rats acquired a larger tendency to sign-track compared to males. However, despite this group difference, at the end of conditioning there was enormous heterogeneity in each group **C)**, with both males and females showing a large number of intermediates, sign- and goal-trackers. **B)** The distribution of subjects across the possible values for index is shown as a box plot.

**Table 1:**
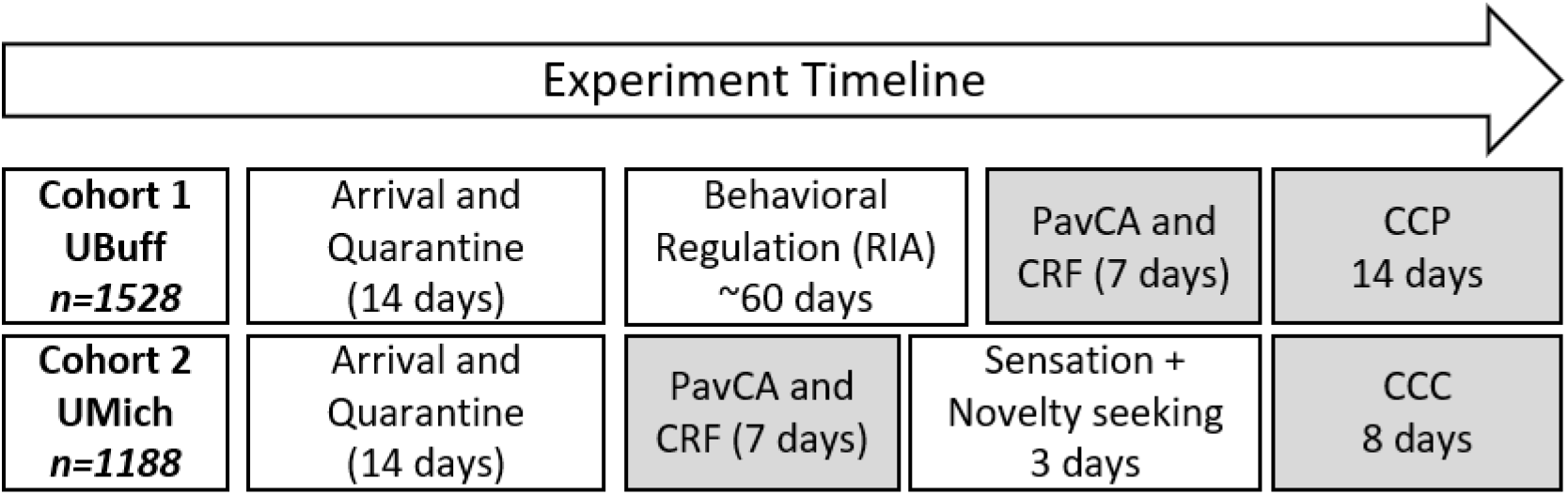
Timeline for the University at Buffalo and Michigan cohorts. Rats arrived at both testing sites, and were quarantined for 14 days before entering the study. Note that the UBuff cohort was tested at the Research Institute on Addictions on several non-drug behavioral regulation tasks before entering testing at the University at Buffalo. Blocks shaded in gray reflect the data presented in this paper.

### Pavlovian Conditioned Approach (UBuff)

During conditioning, rats learned to approach both the lever [main effect of Session: *F*(4,6088)=1026.2; *p* < 0.001], and the food cup [main effect of Session: *F*(4,6088)=44.3, *p* < 0.001] during the 8-second lever-CS period. To quantify individual differences in tendency to goal- and sign-track, the PavCA index was computed yielding a value ranging from −1 (goal-tracking) to 1 (sign-tracking), reflecting the overall tendency to approach the lever or food cup, as described previously:^15^. Relative to males, females showed a greater tendency to sign-track as reflected by a higher terminal index score [Session x Sex: F(4, 6088) = 11.3, *p* < 0.001] **(Fig 1A, B.)**. This is consistent with the findings from the UMich cohort^27^. Despite the observed group sex difference, there was considerable individual variability, with a substantial number of sign- and goal-trackers, as well as intermediates across both sexes **(Fig. 1C)**.

#### Box and Jitter Plots

Because of the large sample size, we have opted to present the data shown in Fig. 1C and elsewhere as box plots **(Fig. 1B)**. The notches reflect the 95% confidence interval for the distribution, where the center of the notch reflects the median. The colored regions of the box plot reflect the inner quartile range, while the remaining outer range of the plot reflects the outer quartile range. The vertical white line with hair ticks reflects the standard error of the mean (SEM), with the center of the line reflecting the location of the mean. Individual subject data is shown behind box plots as a jitter plot. All other box plots presented in this paper follow the same rules for plotting as **Fig. 1B**.

**Fig. 2** shows the time course of acquisition of lever- and food cup-directed responses in intermediates, STs and GTs, during the 5 sessions of PavCA training. As expected, sign-trackers showed higher probability of interaction with the lever [main effect of PavCAPheno: *F*(8,6088)=447.6, (*p* < 0.001)] **(Fig. 2A)**, a larger number of lever deflections [main effect of PavCAPheno: *F*(8,6088)=537.7, (*p* < 0.001)] **(Fig. 2B)**, and a shorter latency to deflect the lever [main effect of PavCAPheno: *F*(8,6088)=447.6, (*p* < 0.001)] **(Fig. 2C)**, compared to goal-trackers. Similarly, goal-trackers showed a higher probability of entering the food-cup [main effect of PavCAPheno: *F*(8,6088)=502.9, (*p* < 0.001)] **(Fig. 2D),** a larger number of food-cup entries [main effect of PavCAPheno: *F*(8,6088)=428.6, (*p* < 0.001)] **(Fig. 2E)**, and a shorter latency to enter the food cup [main effect of PavCAPheno: *F*(8,6088)=480.3, (*p* < 0.001)] **(Fig. 2F)**, compared to sign-trackers. Thus, the ST/GT/IN phenotype distinction was robust across multiple PavCA behaviors. All six of these measures interacted with sex [Session X PavCAPheno X Sex interactions: *F*s(8,6088):= 19.9, 8.4, 19.5, 2.5, 4.1, 3.5, (*p*s < 0.01)], such that female sign-trackers and intermediates showed more lever deflections, quicker latency, and higher probability of lever contact that male sign-trackers and intermediates **(Fig. 2)**. Similarly, female goal-trackers and intermediates showed more food-cup entries, quicker latency, and higher probability of food cup entry and males did **(Fig. 2)**. During the intertrial interval between lever presentations, females also showed a higher tendency to engage the food-cup across all 5 sessions [Sex X Session interaction: *F*(4,5980)=76.2, *p*<0.001] (data not shown), suggesting increased general activity in females relative to males. Again, these results are very similar to those described in the UMich cohort^27^.

**Figure 2:**
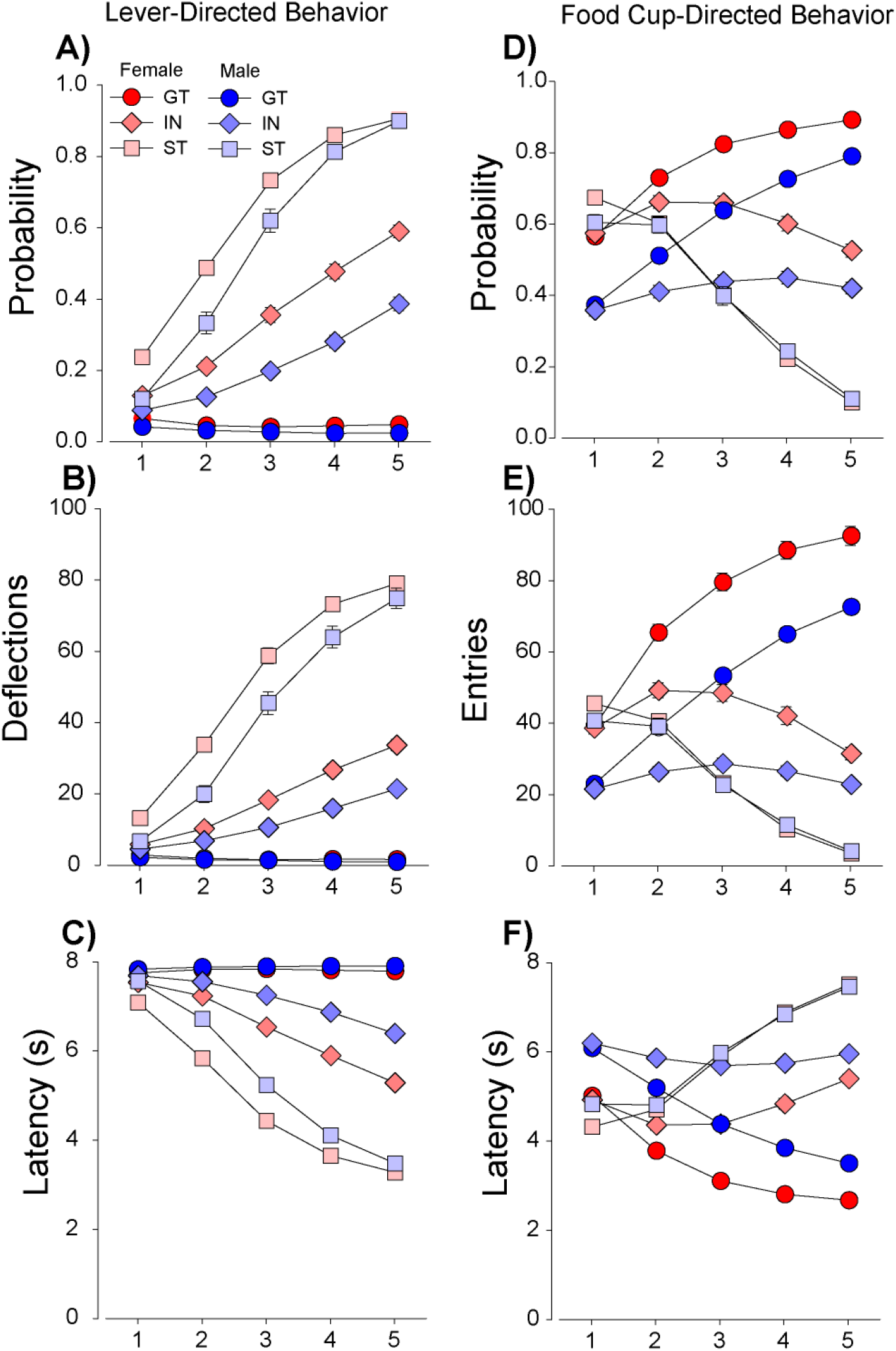
Individual measures of tendency to sign- and goal-track used to calculate PavCA index (UBuff). Performance of intermediates, sign- and goal-trackers across 6 major behavioral measures during Pavlovian conditioned approach. Sign trackers showed **A)** a higher probability of deflecting the lever, **B)** increased number of lever deflections, and **C)** faster latency to deflect the lever than intermediates and goal-trackers. Further, this effect was larger in females across all three PavCA phenotypes. Conversely, goal-trackers showed **D)** higher probability of entering the food cup, **E)** more food cup entries, and **F)** quicker latency to enter the food cup than intermediates and sign-trackers. Similarly, this effect was more robust in females across all three measures.

### Conditioned Reinforcement

Next, rats were tested for the conditioned reinforcing properties of the lever stimulus using a Conditioned Reinforcement (CRF) test, in which rats learned to nosepoke for presentations of the lever-CS, as described previously^28^. During conditioned reinforcement, three measures of the conditioned reinforcing effect of the lever were measured: 1) active-directed responses, 2) number of earned lever presentations, and 3) number of lever deflections per presentation.

The lever served as an effective reinforcer in all rats [main effect of Port: *F*(1,1522)=1718.0, (*p* <0.001)], although it was a more effective conditioned reinforcer in sign-trackers than goal-trackers and intermediates for all three measures: active responding [Port x PavCAPheno: *F*(2,1522)=93.7, (*p* < 0.001)] **(Fig. 3A, shown as active – inactive)**, earned reinforcers [main effect of PavCAPheno: *F*(2,1522)=196.4, (*p*<0.001)] **(Fig. 3B)**, and lever presses per reinforcer [main effect of PavCAPheno: *F*(2,1510)=83.4, (*p*<0.001)] **(Fig. 3C)**. Further, this effect was also stronger in females for both earned reinforcers [PavCAPheno X Sex interaction: *F*(2,1522)=3.5, (*p*<0.05)] and lever deflections per reinforcer [PavCAPheno X Sex interaction: *F*(2,1510)=6.3, (*p*<0.01)]. Indeed, variance in PavCA accounted for 38-48 % of the variance in lever presses per reinforcer in males and females, respectively (*p*<0.05) **(Fig. 3F)**. This particular measure most directly reflects the incentive value of the lever during this task; in comparison the other two measures, while also significantly correlated, accounted for much less of the variability between PavCA and CRF **(Fig. 3D, 3E)**. These results are also consistent with those reported in the UMich cohort^27^, and further support the notion that the lever-CS was attributed with greater incentive salience in sign-trackers than goal-trackers^28^.

**Figure 3:**
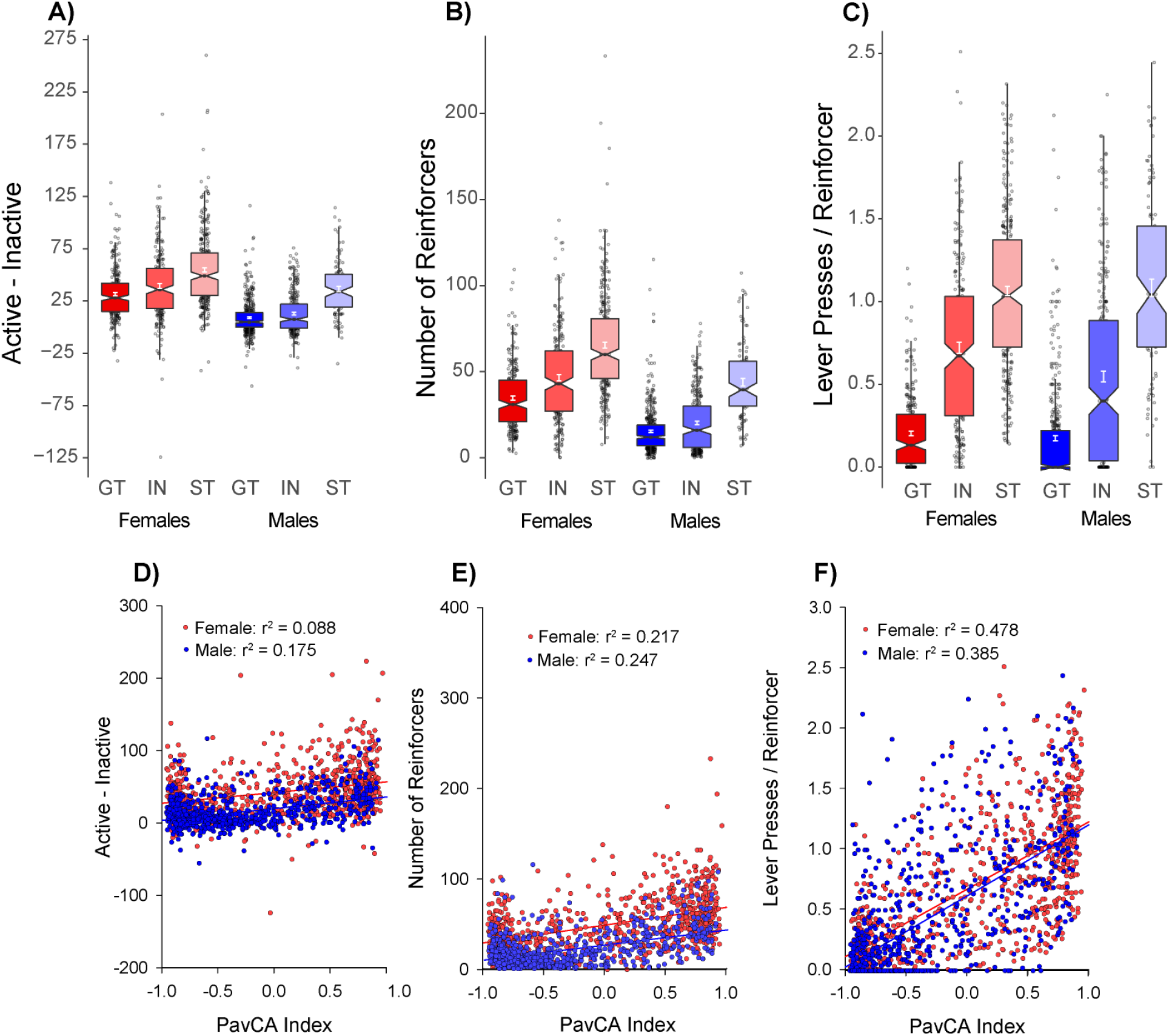
Conditioned Reinforcement (UBuff): Intermediates, sign- and goal-trackers learned to nosepoke for 3-second presentations of the conditioned lever stimulus. All subjects directed their responding to the active hole **A)**, although this effect was largest in sign-trackers. Consequently, **B)** the number of times the subject was reinforced was also larger in sign-trackers than intermediates and goal-trackers. Further, **C)** sign-trackers interacted with the lever stimulus the most during this test, followed by intermediates and goal-trackers. Across all subjects, PavCA index was correlated with **D)** number of responses for the lever-CS, **E)** earned lever-CS reinforcers, and **F)** lever presses per reinforcer. The strongest correlation was between index and lever-directed pressing behavior shown in **F)**, presumably because these two measures are the most strongly related.

### Cocaine Cue Preference: Locomotion

Next, we observed that the relationship between the propensity to attribute incentive salience to a food CS, and the unconditioned locomotor effects of cocaine were independent the UBuff cohort. Specifically, a cocaine conditioned cue preference (CCP) procedure was used, whereby locomotor activation following a 10 mg/kg i.p. injection of cocaine was measured over four conditioning trials. Importantly, each trial consisted of a single injection of cocaine and saline, in alternation, on a discrete cocaine or saline paired floor stimulus. Cocaine induced significant locomotor activation compared to saline across all 4 conditioning trials [Drug x Trial interaction: *F*(3,4572)=14.9, (*p* < 0.001)], which increased by trial 4 compared to trial 1, reflecting sensitization **(Fig. 4C).** Further, the locomotor activating effect was larger in females than in males across each of the 4 sessions [Drug x Sex interaction: *F*(1,1524)=419.9, (*p* < 0.001)] **(Fig. 4C).** However, there was no significant effect of PavCAPheno on either Day 1 or Day 4 of testing (*ps* > 0.05) **(Fig. 4A, 4B)**, indicating that INs, STs and GTs did not differ in their locomotor response to cocaine. Indeed, there was no correlation between the PavCA Index score and cocaine-induced locomotor activity in either males or females **(Fig. 4D)**. When all 4 sessions of conditioning were evaluated in a single analysis, there was a significant Drug X Trial X PavCAPheno interaction [*F*(6,4572)=2.17, (*p* < 0.05)],)], but the effect size was very small (η^2^ = 0.003). Further, although there was an effect PavCA phenotype on baseline locomotion during the habituation session [main effect of PavCAPheno: *F*(2,1522)=10.2, (*p*<0.001)] and on the first saline trial [main effect of PavCAPheno: *F*(2,1522)=6.1, (*p*<0.001)], these effect sizes were also small (η^2^ = 0.013, 0.007 respectively). Thus, tendency to sign- or goal-track is largely unrelated to the locomotor response of both acute and repeated injections of cocaine at the 10 mg/kg dose during CCP, and does not appear to be meaningfully related to locomotion under non-drug conditions.

**Figure 4:**
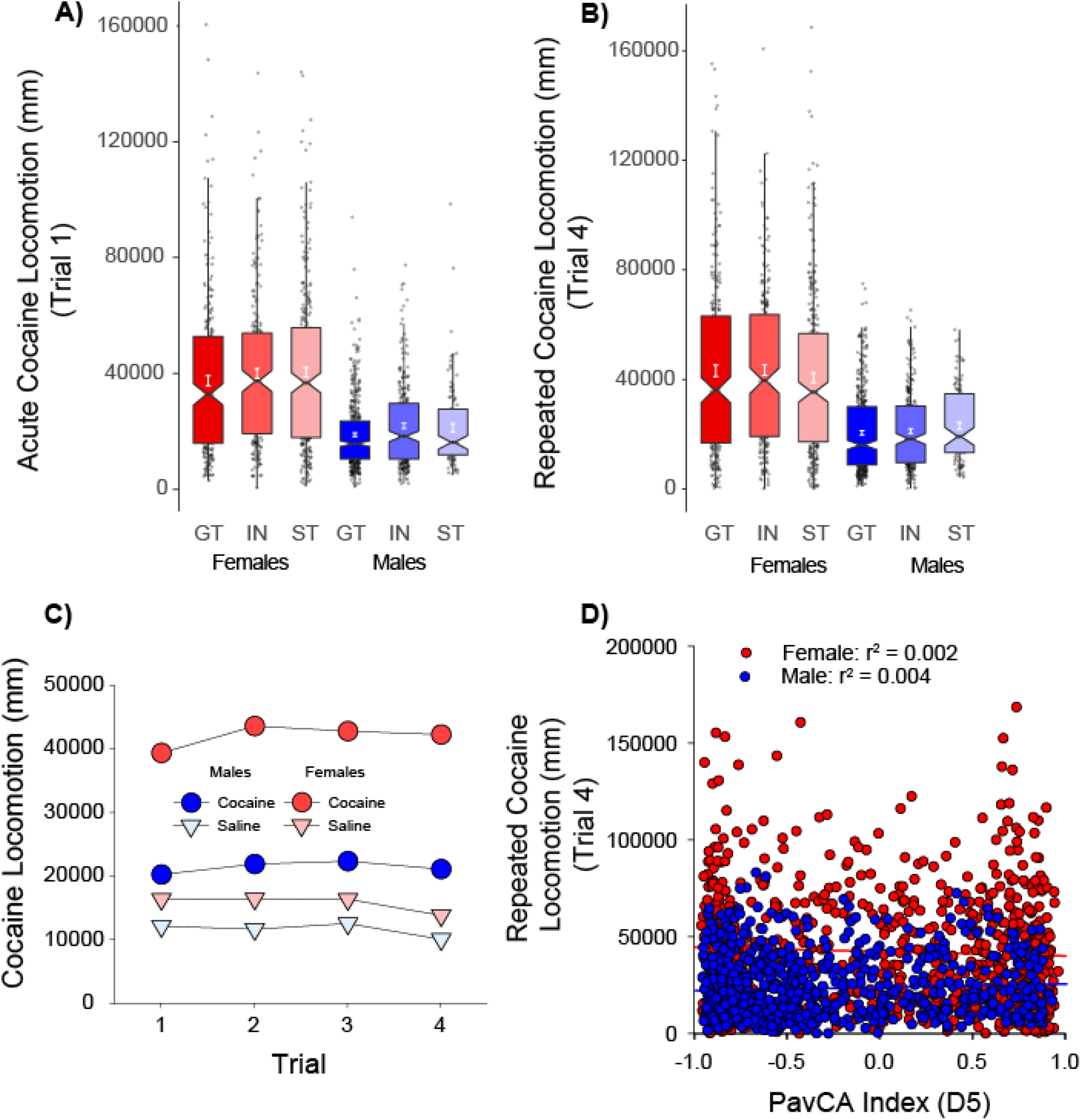
Unconditioned locomotor response during CCP (UBuff): Rats showed robust locomotor activation to cocaine across 4 trails of conditioning. On the first trial **A)** females showed larger cocaine-induced locomotion compared to males, although this effect was not informed by PavCA phenotype. **B)** This sex difference persisted to the fourth trial of conditioning. **C)** Cocaine induced larger locomotor activity compared to saline across trials, and this effect sensitized between sessions 1 and 4. **D)** PavCA index is largely unrelated to cocaine induced locomotion at the end of conditioning in both males and females.

### Cocaine Contextual Conditioning: Locomotion

Although a 10 mg/kg dose of cocaine is on the ascending limb of the cocaine locomotor dose response curve^29^, one of the hallmark features of cocaine sensitization is the development of stereotypy^30^ which include repetitive head movements (head waving) in rodents. Thus, in the UMich cohort of rats, a cocaine contextual conditioning (CCC) procedure was used to examine the development of both-cocaine induced locomotion and bouts of headwaving thought to reflect instances of stereotypy at the start and end of 5 daily conditioning trials to a 15 mg/kg dose of cocaine. In this case, headwaving bouts were computer recorded when the rat was not exhibiting locomotion, but in place and moving its head side-to-side, as previously described^24^.

Unlike CCP, here rats underwent testing in a constant context characterized by a wire-mesh floor and grey walls. Rats were allowed one session to habituate to the testing environment (day 1), followed by a baseline session (day 2), prior to which all rats received saline injections. On the first cocaine conditioning trial (day 3), cocaine acutely increased locomotor activity relative to baseline day 2 [main effect of Trial: *F*(1,1167)=1926.2, (*p* < 0.001)], and this effect was more robust in females [Trial X Sex: *F*(1,1167)=63.4, (*p*<0.001)] **(Fig. 5A)**. We did find a three-way interaction [Trial X PavCAPheno X Sex: *F*(2,1167)=3.1, (*p*<0.05)] which revealed that on the first cocaine treated day (day 3), female STs showed a modest increase in locomotor activity compared to STs, although neither STs or GTs differed from INs. There was no effect of PavCAPheno in males **(Fig. 5A)**. Cocaine also produced modest bouts of head waving on the first trial [main effect of Trial: *F*(1,1167)=83.7, (*p* < 0.001)], and this effect was greater in females than males [Trial X Sex: *F*(1,1167)=56.0, (*p*<0.001)], but was not affected by PavCAPheno (**Fig. 5C)**.

**Figure 5:**
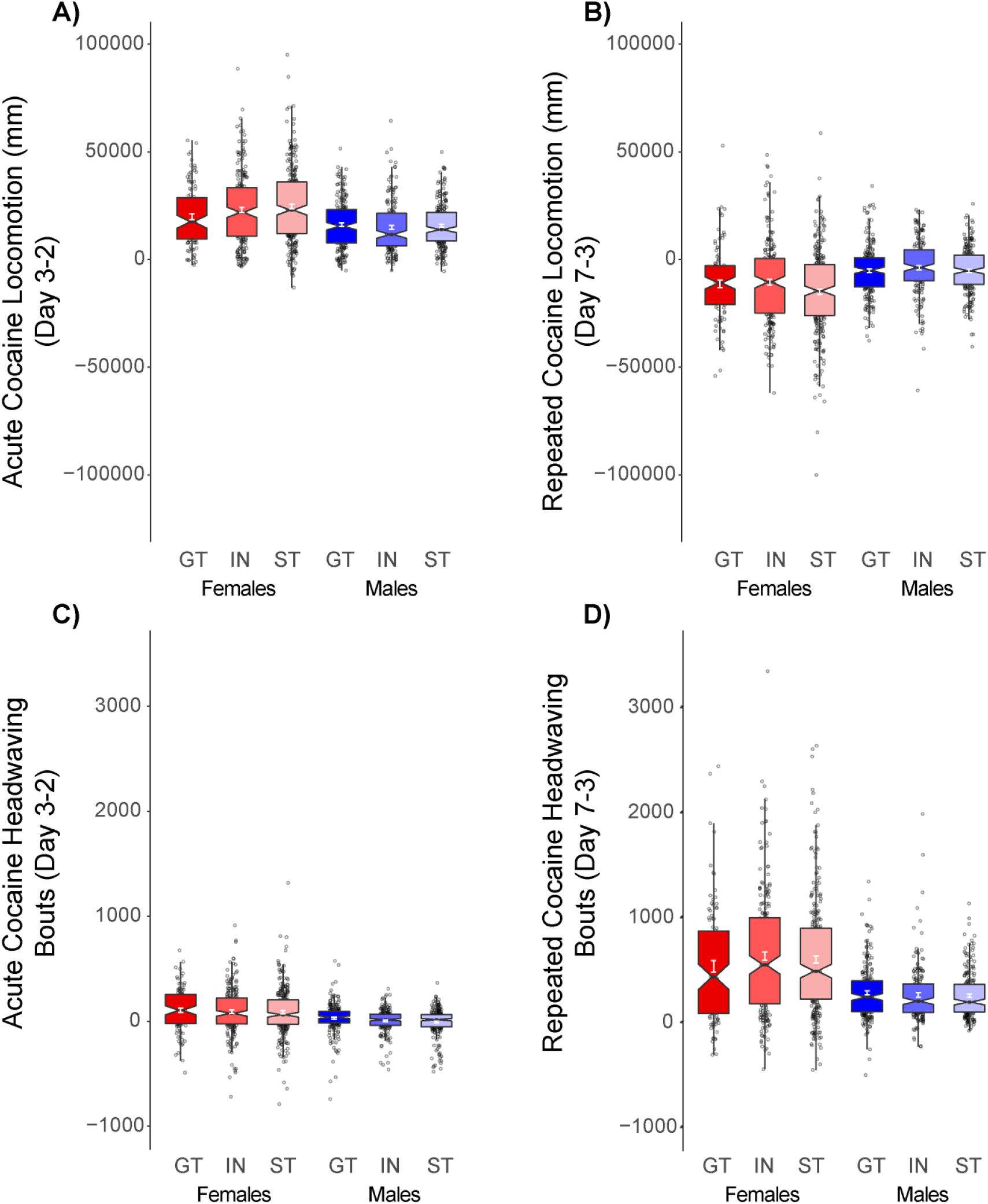
Unconditioned locomotor response during CCC (UMich): Rats showed **A)** significant locomotor activation on the first session of conditioning compared to baseline, and this effect was larger in females than males. Further, **B)** cocaine treatment induced headwaving on this first session as well, although neither locomotor activity or headwaving was related to tendency to sign- or goal-track. On the final day of conditioning, subjects showed a **C)** decrease in cocaine induced locomotion, and **D)** an increase in cocaine-induced headwaving, and this effect was higher in females. Further, this effect was independent from PavCA phenotype.

We next examined the development of sensitization following five days of conditioning (day 3-7). Here, all subjects showed a decrease in locomotor activity by the end of conditioning [main effect of Trial: *F*(1,1167)=276.6, (*p*<0.001)], and this decrease was more pronounced in females than males [Trial X Sex: *F*(1,1167)=58.1, (*p*<0.001)]. That is, females showed the greatest change in locomotor behavior from day 3 to day 7 **(Fig. 5B)**. This decrease in locomotor activity was accompanied by a robust increase in head waving [main effect of Trial: *F*(1,1167)=968.1, (*p*<0.001)], and again this effect was larger in females than males [Trial X Sex: *F*(1,1167)=145.2, (*p*<0.001)] **(Fig. 5D)**. There was, however no effect of PavCAPheno following the 5 conditioning sessions, and tendency to sign- or goal-track appears largely unrelated to the unconditioned locomotor activating effects of cocaine.

### Cocaine Cue Preference: Conditioned Approach

The tendency to sign-track is characterized by approach to reward-predictive cues. To test whether the tendency to sign-track was related to approach a discrete drug-paired cue, in this case a tactile floor, the change in preference for a cocaine-paired floor stimulus during CCP was examined. Subjects in the UBuff cohort were measured for “grid” and “hole” floor preference before and after being paired with 4 injections of 10 mg/kg cocaine. Subjects were counter-conditioned, such that cocaine was paired with the less preferred floor stimulus during the pre-test. Previously, Sprague-Dawley sign-trackers showed a robust conditioned cue preference to a cocaine-paired floor stimulus, whereas goal-trackers did not^3^. Here, we sought to characterize this relationship using the population heterogeneity of HS rats, and although we expected that sign-trackers would show more robust expression of cue preference, here we observed these two traits were unrelated.

All groups showed a significant increase in preference for the cocaine paired floor following conditioning [main effect of Test: *F*(1,1522) =1029.7, (*p*<0.001)], and this effect was larger in females than males [Test X Sex: *F*(1,1522)=8.22, (*p*<0.01)]). Goal-trackers actually showed the greatest increase in preference for the cocaine paired floor [Trial X PavCAPheno: *F*(2,1522)=4.83, (*p*<0.01)] **(Fig. 6A)**. However, this is likely due to the counter-conditioning design used. During the pre-test, goal-trackers exhibited stronger bias against the cocaine paired floor prior to conditioning (data not shown). Further, PavCA index was uncorrelated with time spent on the cocaine-paired floor **(Fig. 6B)**. Whether the PavCA index score was related to an increase in locomotor activity during the post-test relative to pre-test, was also examined, but these two measures were not significantly correlated **(Fig. 6C).** Thus, tendency to sign- or goal-track did not meaningfully inform magnitude of conditioning for the cocaine-paired floor type.

**Figure 6:**
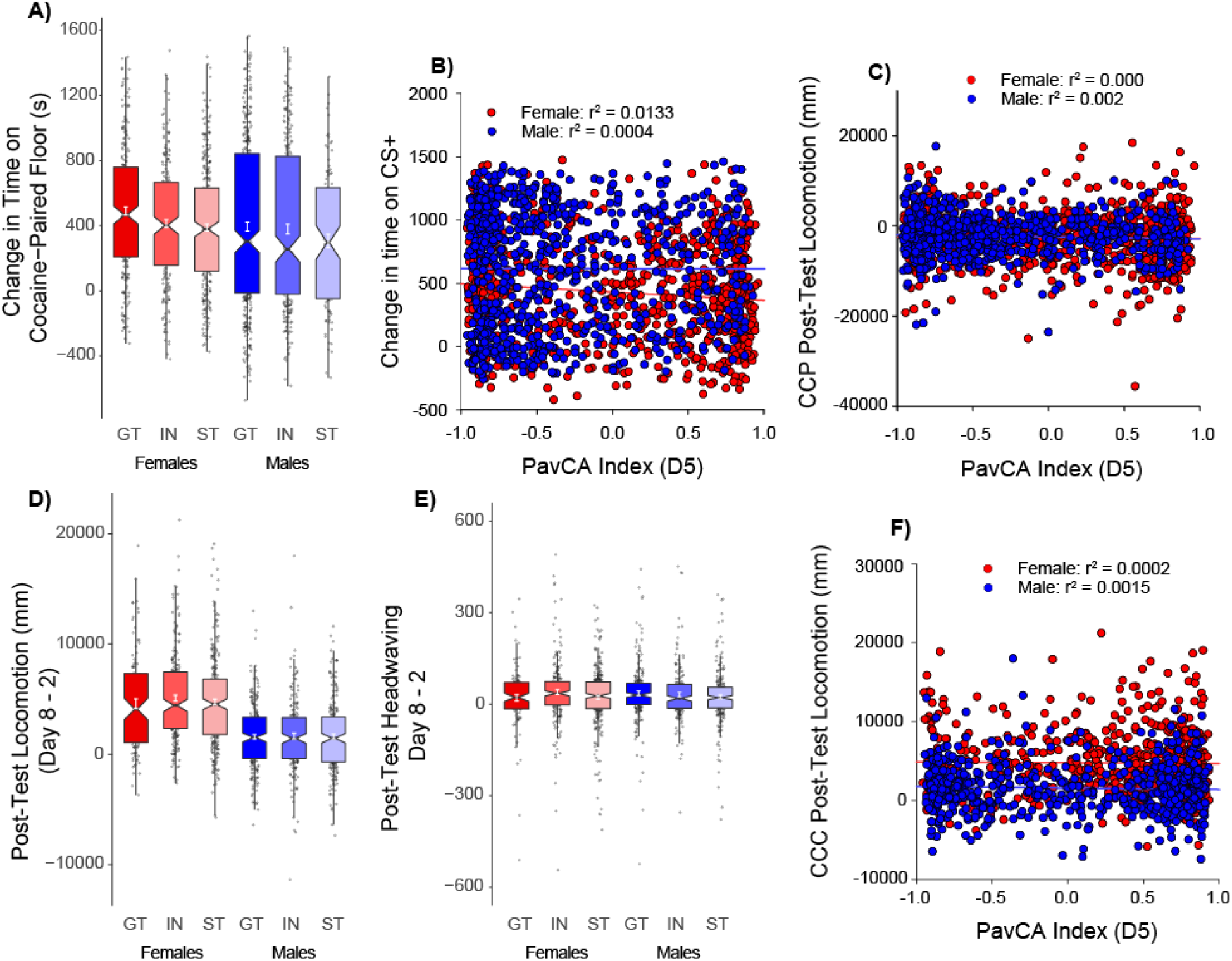
Conditioned approach and locomotion during CCP and CCC (UBuff + UMich): During CCP following 4 saline and cocaine parings with two different tactile floor types, **A)** subjects showed an increase in time spend on the cocaine paired floor following conditioning. However, **B)** despite the heterogeneity in magnitude of conditioning, change in time spent on the cocaine paired floor showed no correlation PavCA index. Further, **C)** locomotor activity instigated on the post-test by the presence of the cocaine-paired floor was also unrelated to PavCA index. During CCC, **D)** subjects showed increased conditioned locomotion on the posttest by the cocaine paired context, and this effect was larger in females. However, neither conditioned locomotion, nor **E)** conditioned headwaving were influenced by PavCA phenotype. Hence, **F)** no significant correlation was detected between index and conditioned locomotor activity.

### Cocaine Contextual Conditioning: Conditioned Locomotion

We further examined the UMich cohort to determine whether a cocaine-paired context would elicit conditioned locomotion following conditioning, and whether this was related to behavior during PavCA. Unlike CCP where a tactile floor was the cocaine predictive stimulus, here the whole testing environment served as a cocaine-predictive context. Locomotor activity during an initial session prior to conditioning (day 2; prior to conditioning, but following saline injection) was compared to that on day 8, which followed 5 sessions (day 3-7) of cocaine injections paired with that context conditioning (day 8). On day 8, exposure to the cocaine-paired context elicited greater locomotor activity than that on day 2 [main effect of Test: *F*(1,1167)=789.4, (*p* < 0.001)], and this effect was larger in females [Test x Sex: *F*(1,1167)=194.5, (*p* < 0.001)] **(Fig. 6D)**. However, there was no effect of PavCAPheno on conditioned locomotion (*p*s > 0.05). Similarly, although there was a modest increase in observed headwaving following conditioning [main effect of Test: *F*(1,1167)=85.7, (*p* < 0.001)], this effect was minimal compared to conditioned locomotor activity **(Fig. 6E)** and was not related to Sex or PavCAPheno. PavCA index was not correlated with conditioned locomotion **(Fig. 6F)**. These results indicate that attribution of incentive salience to reward cues, as measured by approach to a food CS (ST), is independent of both approach to a cocaine-paired floor stimulus, and the conditioned locomotor response to a cocaine-paired context.

### Principal Components Analysis

Principal components analyses were conducted to determine whether the relationship between conditioned and unconditioned responses to cocaine, and the propensity to attribute incentive salience to a food CS could be reduced to fewer dimensions. The measures included the primary measures from PavCA (Index and lever directed behavior during CRF), the acute and repeated unconditioned locomotor responses to cocaine, and conditioned approach and conditioned locomotion responses to cocaine-paired stimuli. When the entire population of both cohorts was included in each analysis, two major factors accounted for the majority of variance in these measures.

PCA for the UBuff PavCA-CCP cohort revealed that two factors accounted for 64% of the total variance. The first factor, which accounted for 36.9% of the variance, had strong loadings from both lever presses per reinforcer during CRF, and terminal PavCA index (> 0.9), with non-significant loadings from CCP measures **(Fig. 7A)**. Conversely, factor 2 had strong loadings from Trial 1 and Trial 4 cocaine induced locomotion (> 0.8), with non-significant loadings from change in time spent on cocaine-paired floor and PavCA measures **(Fig. 7A)**. Together, this further supports the notion that PavCA, and the unconditioned and conditioned measures during CCP are independent.

**Figure 7:**
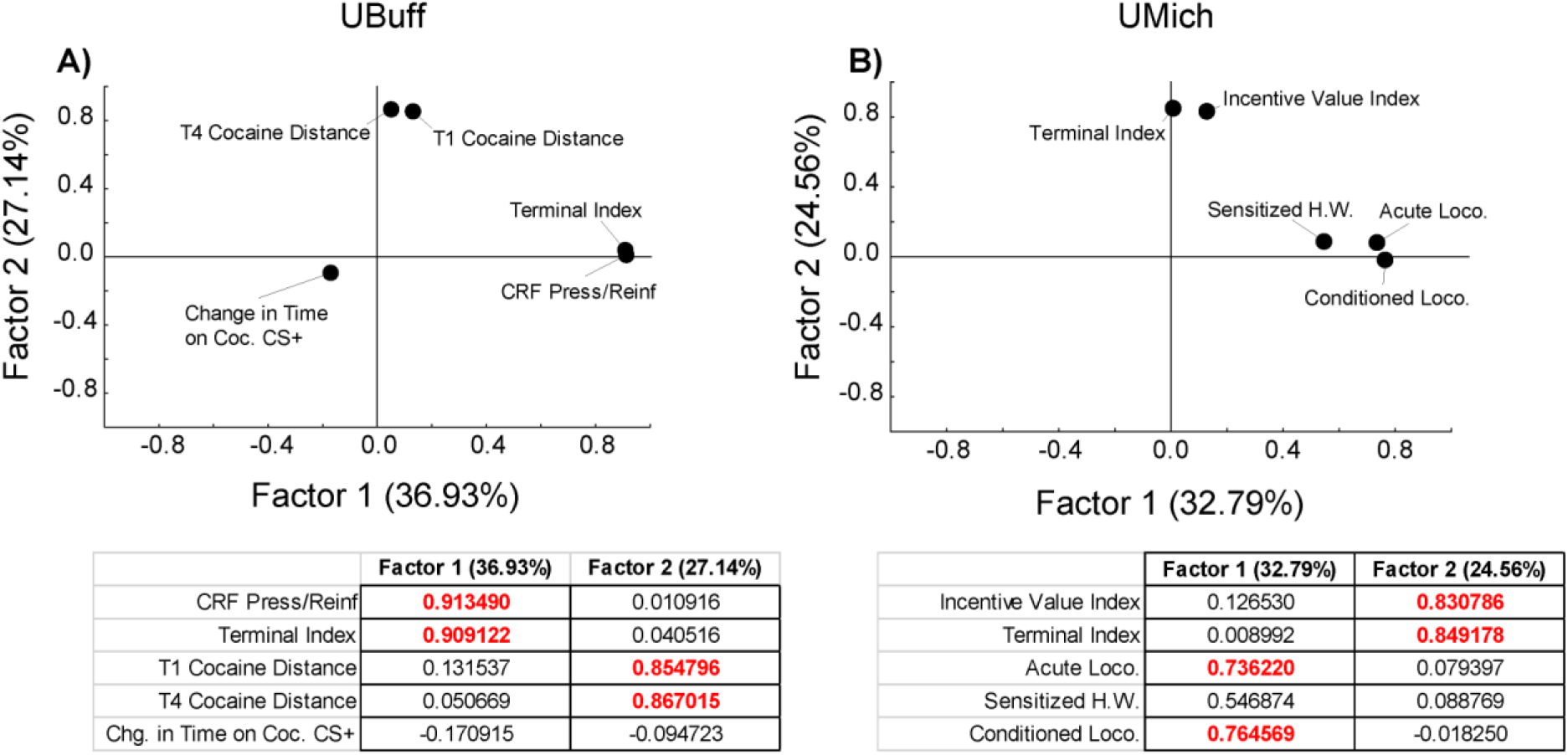
Principal components analysis of PavCA, CCP, and CCC (UBuff + UMich). In both cohorts, we examined whether the traits examined could be reduced to more basic dimensions using principal components analysis. For both cohorts, two factors explained a majority of variance in these studies. In the Buffalo cohort **A)** factor 1 showed significant loading from terminal index and interaction with the lever during CRF, while factor 2 showed significant loading from cocaine-induced locomotion during CCP. Similarly, in the Michigan cohort **B)**, factor 1 showed significant loading from the immediate and conditioned locomotor response to cocaine, whereas, factor 2 showed significant loading from terminal index and incentive value index during CRF. Individual factor loadings for included measures are shown below each panel. Red highlighted values indicate factor loadings that exceed 0.7.

A similar pattern of results was found for the UMich PavCA-CCC cohort, where two factors accounted for more than half the variance. The first factor accounted for ~38% of the variance and contained strong loadings from both acute locomotion and the conditioned locomotor response to the cocaine-paired context (> 0.7) (**Fig. 7B)**, with non-significant loadings from the PavCA measures and sensitization of the headwaving response. Factor 2, by comparison, had strong loadings (>0.8) from both PavCA measures **(Fig. 7B)** and non-significant loadings from the CCC measures. This finding further suggests that, similar to CCP, behavior during CCC and PavCA are largely unrelated to each other.

## Discussion

The purpose of the present study was to elucidate correlations between multiple addiction-related traits in a large sample of genetically diverse heterogeneous stock rats. The advantage of this approach is that is allows for multiple within-subjects comparisons across a broad range of behaviors and testing paradigms, each of which are thought to examine different but arguably related psychological processes. To this end, a total sample size of *n* = 2,701 rats was used to examine Pavlovian conditioned approach and its relationship to both conditioned and unconditioned responses to cocaine in two different tasks. Specifically, two cohorts of rats, one from UBuff and the second from UMich, were phenotyped for their tendency to engage a reward-predictive lever stimulus during the Pavlovian conditioned approach task. Next, performance during PavCA was compared to performance during CRF, and to the locomotor activating effects of cocaine acutely and after repeated exposure during either a CCP or CCC task. Finally, PavCA and CRF was compared to the conditioned approach to a cocaine-paired floor, or conditioned locomotion to a cocaine-paired context during CCP and CCC, respectively. These findings were largely consistent with previous reports. First, the tendency to sign- or goal-track was significantly correlated with performance during the conditioned reinforcement test^15,21,27,28^, in which rats were allowed to nosepoke for presentations of the Pavlovian lever-CS. Second, although there was substantial variability in locomotor activation during both CCP and CCC, PavCA did not correlate with locomotion during either task, consistent with others who have also reported the tendency to sign-track as having either a subtle or unrelated effect on locomotion to a novel environment^31^ or following cocaine treatment^5,18,32^. Indeed, unlike selectively-bred high-responder rats, who show increased locomotion to a novel environment and a greater tendency to sign-track relative to bred low-responders^33,34^, both locomotion and tendency to sign-track were independent in HS rats. Third, whereas food cues acquire incentive motivational properties to a much greater extent in STs than GTs, our hypothesis that STs would prefer a cocaine-associated floor cue (based on ^3^) was not supported. Finally, the conditioned locomotor effects during CCC were not related to PavCA. Thus, there were no meaningful correlations or relationships between performance during PavCA with either CCP or CCC, which ultimately leads us to suggest that these traits are largely independent of one another.

Previously, Pavlovian conditioned approach has been associated with performance on variety of other traits, including drug-conditioning^1,2,18,19,35–37^ and non-drug behaviors^21,22^. At least in HS rats, the relationship between tendency to sign-track and a subset of these other traits may be dissociable. In support of this notion, it has recently been demonstrated that PavCA performance is independent of sensation- and novelty-seeking in a large sample of HS rats^27^, suggesting drug-related traits can be dissociated in a sufficiently diverse and large subject pool. This is the first instance in which the HS strain been used extensively for cocaine conditioning in relation to PavCA, and independence of these particular traits may reflect genotypic and phenotypic diversity unique to the HS population that is not present in other commonly used strains such as Sprague-Dawley rats. Not every behavioral task was independent from each other, as PavCA was related to CRF, suggesting that there is a fundamental dissociation between the processes underlying attribution of incentive salience to reward cues, CCP, and CCC within HS rats. Our PCA factor loadings further support this finding, in that measures of incentive salience (terminal index and lever-directed behavior during CRF) showed independent loadings from conditioned and unconditioned locomotor activation in both cohorts of rats. Although other populations of rats may indeed show a different degree of relatedness between these particular tasks, HS rats in particular would be useful in examining each of these particular traits in isolation of the other.

In addition, there were pronounced sex differences across each of the tests employed here, particularly during PavCA, CRF, and the locomotor effects of cocaine in CCP and CCC. While we are not the first to report sex differences during PavCA^21,27,38,39^, cocaine-induced locomotion^40,41^, or place preference^42^, we have replicated a variety of previous reports on these differences using a large sample size. In light that these traits are arguably independent, the replication of previously identified sex differences reinforces the need to examine both males and females in the behavioral research, with particular attention to how the biological etiology of behavior differs between the two sexes. The HS line of rats in particular may be a useful tool specifically for examining the genetic and etiological basis of sex differences within a specific behavior of interest, while being able to dissociate it from other previously linked traits.

This study is the first to examine unconditioned and conditioned cocaine responses in relation to Pavlovian conditioned approach with a large sample size. Most of the behavioral measures in both tasks were largely unrelated, and those related traits that were identified showed marginal contribution to the unconditioned effects of cocaine. These data underscore the importance of conceptualizing addiction more generally as a multifaceted process, in which multiple independent traits and pathways may result in maladaptive drug use behavior. Further, we characterized robust sex differences across each of the behavioral paradigms used here. In summary, HS rats may serve as useful tool for examining a specific trait independently of others. Further, although these traits may be similar in other strains, caution should be used in interpreting results across studies using different subjects and sample sizes.

## Materials and Methods

### Subjects

NMcwi:HS (here after referred to as HS) rats were shipped from the laboratory of Dr. Leah Solberg-Woods at Wake Forest University School of Medicine to either the Department of Psychology at the University at Buffalo (UBuff; n = 1528) or the Department of Psychology at the University of Michigan (UMich; n = 1188) at approximately 33 days old, as part of the NIDA Center for GWAS in Outbred Rats. These HS rats were established at the NIH from eight founder strains of separate lineages^43^, and are maintained using 64 breeding pairs using a breeding scheme that accounts for kinship coefficients to minimize inbreeding and maintain genetic heterogeneity. They show high genotypic and phenotypic diversity, and are useful for the complex mapping of genetic correlates for a variety of behaviors^44–46^.

Rats of the same sex were pair-housed (UBuff) or triple-housed (UMich) in plastic cages (42.5×22.5×19.25 cm). Cages were lined with bedding (Aspen Shavings) and kept in a temperature-controlled environment (22±1°C). No environmental enrichment was provided throughout the experiment. Water and food (Harlan Teklad Laboratory Diet #8604, Harlan Inc., Indianapolis, IN, USA) were available ad lib and housing was maintained on a 12h reverse light/dark cycle (lights off at 0730h). All testing occurred during the dark phase at least 1 hour following lights off. For rats tested at the UMich, behavioral testing began at approximately 60 days old. Rats were then tested for Pavlovian conditioned approach, “novelty-seeking”, “sensation-seeking”, and cocaine contextual seeking (CCC) as described below and in **Table 1.**

For rats tested at the UBuff, rats first arrived at the Laboratory Animal Facility and were kept in quarantine for 14 days. They were then transferred to the Research Institute on Addictions in Buffalo, NY, and tested in several paradigms at age PND72. These paradigms, the data from which are the subject of a separate publication, included open field locomotion, light reinforcement, choice reaction time task, delay discounting, and social reinforcement. At the beginning of testing, the average weight of females was 197g, and the average weight of males was 315g. The rats were then transferred to the Psychology department and began testing (mean PND162, range 140-204) for Pavlovian Conditioned Approach (PavCA) and Cocaine Cue Preference (CCP) as described below and in **Table 1**. Rats were tested in 16 batches, each batch consisted of 7 groups of 16 subjects. Rats were tested in the same order daily. All studies were conducted according to the National Research Council (2003) “Guidelines for the Care and Use of Mammals in Neuroscience and Behavioral Research”. All procedures were approved by the UBuff and UMich Institutional Animal Care and Use Committees.

### Drugs

During CCP and CCC, subjects were treated with either 0.9% physiological saline, or 10 and 15mg/kg injections (i.p.) of cocaine HCl (Nat. Inst. Of Drug Abuse, Bethesda, MD) dissolved into sterile saline at 10 or 15mg/mL respectively. All injections were given immediately prior to conditioning sessions.

### Apparatus

#### Pavlovian Conditioned Approach (PavCA)

Testing occurred in 16 modular testing chambers (20.5×24.1 cm floor area, 29.2 cm high; MED-Associates Inc., St. Albans, VT) located inside either Med-Associates (UMich) or custom-built (UBuff) sound and light attenuating chambers equipped with fans for ventilation and noise masking (A&B Display Systems, Bay City, MI). 45 mg banana pellets were delivered via a pellet dispenser into a food cup equipped with an infrared photobeam detector to detect head entries. Each chamber contained a retractable backlit lever (2 cm length, 6 cm above floor) on either the left or right side of the food cup. A red houselight was located on top of opposing wall of the chamber (27cm high). During the conditioned reinforcement test, the retractable lever was moved to the center of the wall and the food-cup was removed. Two nosepoke ports with head-entry detectors were situated on the left and right side of the lever. All data were collected using the Med-PC IV software package.

#### Cocaine Conditioned Cue Preference (CCP)

Rats were tested in the dark in black acrylic chambers (47 cm length x 19 cm width x 30 cm height) with either “grid” or “hole” textured floors that were spray painted black. Beneath the textured floors was an additional smooth black matte floor. During testing, subjects were video recorded using infrared cameras connected to a 16-channel DVR (Swann Communications, Inc., Santa Fe Springs, CA) to analyze locomotor activity and side/floor preference. Videos were analyzed in real-time using Topscan video tracking software (Clever Sys. Inc., Reston, VA)^3,24^. All testing environments were located in custom-built light- and sound-attenuating chambers.

#### Cocaine Contextual Conditioning (CCC)

Rats were tested in chambers composed of an outer box (27 in length x 13 in width x 26 in height) and a smaller, insert box (18 in length x 6 in width x 22 in height) that was placed in the center of the outer box. Both the outer box and insert were composed of 4 fiberboard walls and did not include a floor or ceiling. A wire mesh was suspended within the outer box, two inches up from the bottom. The mesh support and the inside of the insert box were painted matte grey with Rust-Oleum automobile primer in order to provide the best contrast for the various colors of rats. Each session of the CCC procedure was video recorded with a Zmodo, ZMD-DR SFN6 DVR and analyzed using Noldus Ethovision motion tracking software.

### Procedure

#### Pavlovian Conditioned Approach (PavCA)

During PavCA, subjects learned the association between presentation of a banana-flavored food pellet and a backlit lever-CS over 5 sessions. In the two days prior to testing, subjects received home cage exposure to banana flavored food pellets (~25 pellets per day; Bio-Serv, Flemington, NJ, #F0059). Rats then received a single day of food cup training to habituate subjects to the testing environment. During food-cup training, subjects underwent a 5-minute chamber habituation period during which the houselight was extinguished. Next, the houselight was illuminated, and subjects received 25 pellets delivered into the food-cup on a VI-30s (1-60s range) schedule. The session ended after the 25 pellets were delivered.

Next, over five daily conditioning sessions, there were 25 lever-food pairings such that delivery of each pellet into the food-cup was preceded by insertion of the lever for 8-seconds. Lever presses had no programmed consequences. Intervals between trials were determined using a VI90 schedule (30-150s range) such that sessions lasted an average of 37.5 minutes.

#### Measures

Across the five conditioning sessions, two conditioned responses were measured during the presentation of the lever-CS: lever-directed approach (number of lever deflections) and goal-directed approach (entries into the food cup). Approach latency and magazine entries during the inter-trial interval (outside of the CS period) were also collected.

Previously, we have used these measures to calculate a **PavCA index;** the general tendency to approach either the lever (“sign-tracking”) or food-cup (“goal-tracking”)^15^. The index is computed by first measuring: 1) The probability differential of contact with the lever versus food-cup during each CS period (average probability of a lever press on a given CS trial – average probability of a food-cup entry on a given CS trial), 2) the response bias directed towards either the lever or the food cup (# lever contacts - # food-cup contacts / # lever + # food-cup contacts), and finally 3) the average latency across trials to initiate contact with either the lever or food-cup (food-cup latency – lever latency / 8). These three measures were averaged together, producing an overall PavCA index between −1 and 1, used to categorize subjects as sign-trackers (STs) and goal-trackers (GTs) based on the average index from sessions 4 and 5 of training^15^. Finally, subjects were classed as goal-trackers if their score was between −1 and −0.5, as intermediates (IN) between −0.5 and 0.5, and as sign-trackers between 0.5 and +1.

#### Conditioned Reinforcement (CRF) test and measures

The ability of the food-associated lever-CS to reinforce the acquisition of a new instrumental response (nosepoking) was assessed the day after Pavlovian conditioning ended. Testing occurred in the same chamber used for PavCA but the center food-cup was removed and replaced with the illuminated backlit lever-CS. On both the left and the right side of the lever-CS were two nosepoke ports, one active and one inactive. All other aspects of the testing environment were identical. Nosepokes into the active hole resulted in insertion of the lever-CS into the chamber for 3s, during which lever deflections were recorded. Nosepokes into the inactive port had no programmed consequences. Sessions lasted 40 minutes. The primary measures were entries into active and inactive ports, lever deflections, and number of earned lever presentations. We chose to separately examine nosepokes from earned lever presentations and lever deflections with the idea that responses into a novel nosepoke port and lever-directed responses might be differ in strength to PavCA Phenotype^27^. We ultimately measured lever-directed behavior by using lever presses made per earned reinforcer, a similar outcome measure used in the UMich cohort, the Incentive Value Index ((responses in active port - responses in inactive port) + number of lever presses).

#### Cocaine Conditioned Cue Preference (CCP)

During CCP, rats learned the association between a textured floor stimulus and a 10 mg/kg injection of cocaine. Throughout the 11 days of testing, rats were weighed daily and placed into individual transport containers (Sterilite Corporation, Townsend, MA), for 15 minutes before being moved into the testing room. Subjects first received one day of habituation in which subjects were injected with saline and then placed in the testing environment with a matte floor for 30 minutes to allow rats to acclimate to the chamber. On the following day, subjects received a saline injection and underwent a 30 minute “pre-test” in which the testing chamber was outfitted with both the “hole” and “grid” floor halves, counterbalanced for left/right position. Subjects were counter-conditioned, such that the least preferred floor (the floor each subject spent the least amount of time on) was assigned as the cocaine-paired floor. On the following 8 days, subjects received 10 mg/kg cocaine and i.p. saline injections (order counterbalanced) on alternating days before being placed in the chamber containing a single floor type. Each pair of cocaine and saline conditioning sessions was termed a trial, for a total of 4 trials. Finally, subjects received a post-test, in which they were tested in the presence of both floors following a saline injection. The time spent on the cocaine-paired floor was measured in seconds on both the pre- and post-test sessions. The change in time spent on the cocaine floor was determined by subtracting post-test time from pre-test time. Distance travelled in mm across all testing days was determined by using Topscan’s locomotor analysis.

#### Cocaine Contextual Conditioning (CCC)

During CCC, rats learned to associate a context with a 15 mg/kg cocaine injection. Importantly, CCC differed from CCP in that cocaine pairings occurred with the entire testing context, rather than with exchangeable tactile floor stimuli. First, subjects underwent a single 30-minute session of exposure to the testing apparatus with no prior injection to measure locomotor response to novelty (Day 1). On the next day (Day 2), subjects received an injection of saline and underwent an additional 30-minute pre-conditioning session. Subjects then began cocaine contextual conditioning (Days 3-7) in which rats were treated with cocaine immediately prior to each test session. Finally, on the last day (Day 8) subjects received a 30-minute post-conditioning test session following an injection of saline, to assess the degree of context conditioned hyperactivity (<u>adapted from: ^47^</u>). Throughout testing, two measures of locomotor activity were computer scored: overall distance travelled, and bouts of head waving as described in^24^. Acute locomotor activation by cocaine was analyzed by comparing Day 3 to Day 2. Locomotor sensitization to cocaine was analyzed by comparing Day 3 to Day 7. The conditioned locomotor response to the cocaine-paired environment was analyzed by comparing Day 2 to Day 8.

### Analyses

Repeated-measures analysis of variance (ANOVA), in conjunction with Tukey’s HSD post-hoc tests, were used to probe significant main effects and interactions. For PavCA, CRF, cocaine CCP, and CCC, Sex (male, female), and PavCAPheno (ST, IN, GT) were the between-subjects factors. For PavCA, Session (1-5) was the within-subjects repeated measures factor. For CRF, Port (active, inactive) was the within-subjects repeated measures factor. Lever-directed behavior during CRF (lever presses, lever presses per reinforcer) were both analyzed separately from nosepoking behavior. For cocaine CCP, conditioning Trial (1-4), Test (pre, post) and Drug (saline, cocaine) were the within-subjects repeated measures factors. For CCC, Trial (1, 5) and Test (Pre, Post) were the within-subjects repeated measures factors. For CCC, of the 1188 phenotyped subjects, 15 were dropped due to data collection error and were casewise excluded from all CCC analyses. Further note that the rats presented in the CCC experiment were used in a separate publication examining the relationship between PavCA, response to novelty, and sensation seeking^27^. For brevity, we do not present a dedicated results section for this particular batch of Pavlovian conditioned approach, as the results of these data are similar to the University at Buffalo cohort, and have been described in detail elsewhere.

Further, because subjects in the UBuff cohort arrived at different ages, we ran all analyses presented here using age at the start of testing as a continuous predictor for each variable. While several dependent variables (food cup CS entries, food cup CS entry probability, food cup CS latency, PavCA index, port responses during CRF, earned reinforcers during CRF, change in time on cocaine CS+, CCP locomotor activity, conditioned locomotion) yielded significant main effects or interactions with age at the start of testing, the effect size of these findings were extremely small. Locomotor activity and conditioned locomotion during CCP had the largest effects of age (η^2^ = 0.054, 0.015), with all other measures yielding η^2^ below 0.005. We therefore excluded age as a factor from the primary findings.

In addition, to determine whether the traits discussed here could be reduced to fewer dimensions, two iterations of principal components analysis were conducted in both populations of animals. Specifically, each analysis included index during Pavlovian conditioned approach, lever directed behavior during CRF (lever presses per reinforcer, incentive value index), the first and last days of locomotor activation during CCP and CCC, and the conditioned approach and conditioned locomotion to the cocaine paired floor and context, respectively. All factors examined were determined with a minimum eigenvalue of 1, and were factor rotated using normalized Varimax. All statistics for all experiments were computed using Statistica 13 (Dell Inc., Tulsa, OK). Box and jitter plots were constructed in R (R version 3.6.1., R Studio, Boston, MA) using the ggplot2 package. Principal components plots were generated in Statistica 13. All plots both were modified to improve visual clarity using Adobe Illustrator 2020 (Adobe, San Jose, CA).

## Data Availability

The datasets generated and/or analyzed during the current study are available on our project webpage (ratgenes.org) or from the corresponding author on reasonable request.

